# Ensemble epistasis: thermodynamic origins of non-additivity between mutations

**DOI:** 10.1101/2020.10.14.339671

**Authors:** Anneliese J. Morrison, Daria R. Wonderlick, Michael J. Harms

## Abstract

Non-additivity between mutations—epistasis—profoundly shapes evolution. It can be difficult to understand its mechanistic origins. Here we show that “ensemble epistasis” is likely a universal feature of macromolecules. Using a simple analytical model, we found that epistasis arises when two conditions are met: 1) a macro-molecule populates at least three structures and 2) mutations have differential effects on a least two of the inactive structures. To explore the relative magnitude of ensemble epistasis, we performed a virtual deep-mutational scan of the allosteric *Ca*^2+^ signaling protein S100A4. We found that 27% of mutation pairs gave ensemble epistasis with a magnitude on the order of thermal fluctuations, 1 kT. We observed many forms of epistasis: magnitude, sign, and reciprocal sign epistasis. Depending on the effector concentration, the same mutation pair could even exhibit different forms of epistasis. The ubiquity of ensembles in biology and its pervasiveness in our dataset suggests that ensemble epistasis may be a universal mechanism of epistasis.

**Significance statement:** Addressing the mechanistic origins of evolutionary unpredictability is critical to understanding how mutations combine to determine phenotype. Here we lay the theoretical foundations and investigate the plausibility of a potentially universal mechanism of unpredictability in macromolecules. Macromolecules often adopt a set of interchanging structures, called a thermodynamic ensemble. Mutations can change the relative population of each structure, introducing unpredictability in the mapping between genotype and phenotype. The conditions under which we expect this to arise are common in macromolecules, suggesting that this form of unpredictability may be pervasive in evolution. We conclude that the thermodynamic ensemble bakes unpredictability into biology and that future attempts to address it might incorporate this mechanistic insight.

## Introduction

Epistasis—when the effects of two or more mutations do not sum—is a common feature of biology. It can be thought of in terms of predictability: to make accurate predictions we must know the effect of a mutation in all genetic backgrounds of interest. Epistasis can hint at biological mechanism [1–6], profoundly shape evolution [7–10], and complicate bioengineering that involves simultaneously introducing multiple mutations [11–13]. It is therefore important to understand the general mechanisms by which epistasis can arise. Such understanding will help us better understand biological systems, explain historical evolutionary trajectories, and improve models to predict the combined effects of mutations.

One important class of epistasis is that which occurs between mutations within a single protein. Some-times, such epistasis can be understood intuitively. In Fig 1A, epistasis arises because the positive charge of mutation *A* is adjacent to the negative charge of mutation *B*. Epistasis arises from an electrostatic interaction between charged residues. Sometimes, however, epistasis can be difficult to rationalize. Fig 1B shows epistasis between two positions distant in the structure. Where does such epistasis come from? Can it be predicted from an understanding of protein biochemistry? Such long distance interactions, as well as high-order interactions involving three or more mutations, can be difficult to rationalize mechanistically.

**Fig 1:**
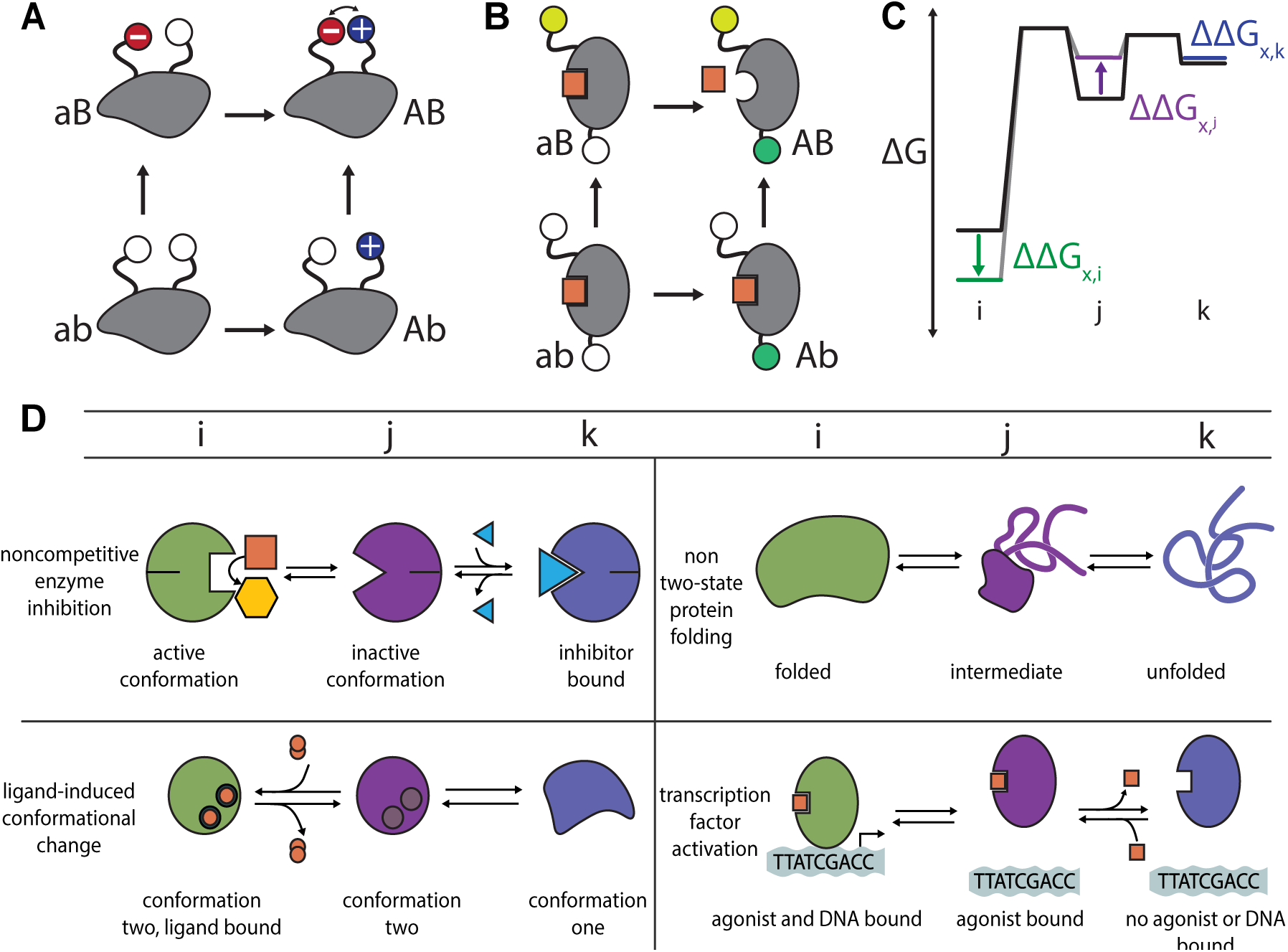
Epistasis and ensembles are common. A) A mutant cycle where epistasis is easily rationalized: the *a* → *A* and *b* → *B* mutations lead to a new electrostatic interaction when introduced together (minus and plus signs) that could explain a non-additive effect. B) A mutant cycle with difficult-to-understand epistasis. Mutations at two distant sites (green and yellow spheres) have no effect on binding of the orange square when introduced independently, but disrupt binding when introduced together. C) Energy diagram for a hypothetical protein that adopts three distinct structures: *i* (green), *j* (purple) and *k* (blue). The y-axis shows the free energy of each structure. The colored arrows indicate the change in the free energy of each structure due to mutation *x*. D) Schematic examples of biological mechanisms in which a protein populates at least three structures. Columns indicate structure labels—*i* (green), *j* (purple), or *k* (blue).

Using a toy thermodynamic model, we argued previously that *ensemble epistasis* could potentially explain such non-additive interactions between mutations [14]. Proteins exist as ensembles of interchanging structures, with the functional output of the protein determined by the average of all structures in the ensemble [15–17]. We found that a mutation can have different effects on different structures in the ensemble, redistributing their relative probabilities in a nonlinear fashion. As a result, the effects of mutations would not sum linearly, leading to epistasis.

Many important questions about ensemble epistasis remain unanswered. Under what conditions is ensemble epistasis expected to arise? Can it lead to different classes of evolutionarily-relevant epistasis, i.e. magnitude, sign, reciprocal-sign, and high-order? Is it plausible that such epistasis could occur in a real protein, rather than the highly simplified lattice models we used previously? And, finally, are there signals for ensemble epistasis that one might detect experimentally?

To address these questions, we set out to rigorously describe the thermodynamic and mechanistic basis for ensemble epistasis. We find that ensemble epistasis arises when a protein populates three or more structures and when mutations have differential effects on structures within the ensemble. This can lead to many types of epistasis, including magnitude, sign, reciprocal sign, and high-order epistasis. Using structure-based calculations on the allosteric S100A4 protein, we predict that a large fraction of mutant pairs could exhibit ensemble epistasis with a magnitude on the order of thermal fluctuations, 1 kT. We also found that varying the concentration of allosteric effectors could tune epistasis, suggesting one might experimentally detect ensemble epistasis by measuring epistasis at different concentrations of allosteric effectors. We conclude that ensemble epistasis is likely an important determinant of non-additivity between mutations in proteins.

## Results

We will treat epistasis as the quantitative difference between the individual effects of mutations *A* and *B* and their combined effect on some quantitative protein property *P* [18]:

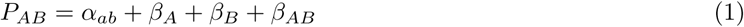

where *P*_*AB*_ is the property of the *AB* genotype, *α*_*ab*_ is the property of the *ab* genotype, *β*_*A*_ and *β*_*B*_ are the individual effects of mutations *A* and *B*, and *β*_*AB*_ is the epistatic term. If *β*_*AB*_ = 0, the effects of *A* and *B* are independent: by measuring *α*_*ab*_, *β*_*A*_ and *β*_*B*_, one can predict *P*_*AB*_ with perfect accuracy. In contrast, for *β*_*AB*_ ≠ 0, the effect of the *A* and *B* mutations depend on the presence or absence of one another. One cannot predict *P*_*AB*_ from the individual effects of *A* and *B* without knowledge of how they interact.

### Mutations affect multiple structures in thermodynamic ensembles

We next constructed a simple model of a protein exchanging between three structures. Our property of interest is the relative population, at equilibrium, of an observable structure *i* relative to structures *j* and *k* (Fig 1C). For example, the population of *i* could quantify the amount of active enzyme in the presence of a non-competitive inhibitor (Fig 1D). This is a generic model that is not subject to the same constraints as the lattice models used previously. It describes, in broad strokes, a wide variety of functions that depend on conformational change (Fig 1D).

The favorability of structure *i* is quantified by the free energy difference between structure *i* and structures *j* and *k*:

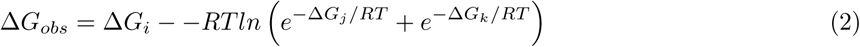

where Δ*G*_*n*_ is the free energy of structure *n* relative to some reference structure, *R* is the gas constant, and *T* is the temperature. Δ*G*_*i*_ describes the free energy of structure *i*, while 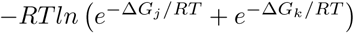 is the Boltzmann-weighted average free energy of structures *j* and *k*. For clarity, we will write this second term as:

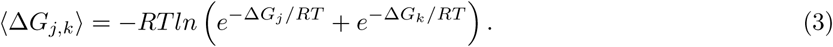

We next considered the effects of mutations on Δ*G*_*obs*_. If a mutation perturbs the physical interactions in a structure, it will change its relative free energy. Because each structure may have different physical interactions, the same mutation may have different effects on different structures. For a three-structure ensemble, we therefore need three terms to describe the effect of mutation *x*: ΔΔ*G*_*x,i*_, ΔΔ*G*_*x,j*_ and ΔΔ*G*_*x,k*_. In Fig 1C, for example, we show a hypothetical mutation that stabilizes *i*, destabilizes *j*, and has no effect on *k*.

To relate this thermodynamic ensemble to epistasis, we need to link changes in genotype to changes in Δ*G*_*obs*_. For bookkeeping purposes, we indicate genotypes as superscripts on each Δ*G* or ΔΔ*G* value. For example, 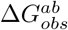 is the observed free energy of genotype *ab*, while 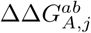 is the effect of mutation *A* on structure *j* in the *ab* background. We assume that mutations are additive within each structure. (By this we mean that 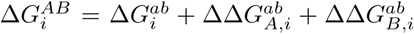. There is no epistatic contribution 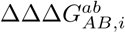 reflecting intramolecular interaction energies within structure *i* of the sort seen in Fig 1A). Using this framework, we can describe the combined effects of mutations *A* and *B* on Δ*G*_*obs*_ as the following:

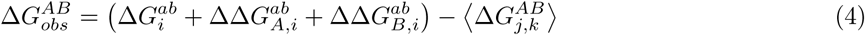

where

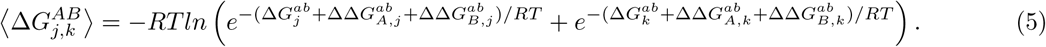

### The map between the thermodynamic and genetic models

To understand the nature of epistasis arising from such a system, we mapped the epistasis model (Equation 1) to the thermodynamic model (Equation 4). This is shown in Table 1.

**Table 1:**
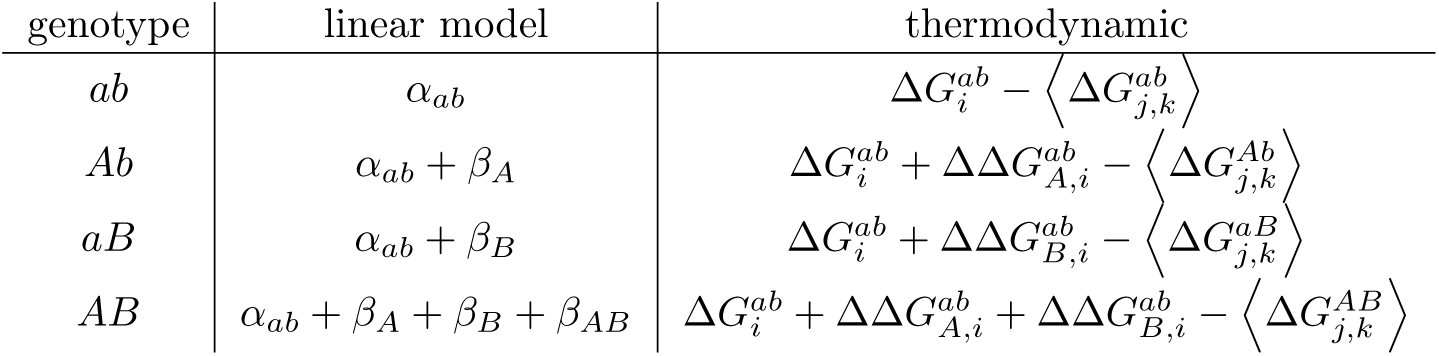
Map between the genetic and thermodynamic descriptions of the observable (Δ*G*_*obs*_).

| genotype | linear model | thermodynamic |
| --- | --- | --- |
| $ab$ | $\alpha_{ab}$ | $\Delta G_i^{ab} - \langle \Delta G_{j,k}^{ab} \rangle$ |
| $Ab$ | $\alpha_{ab} + \beta_A$ | $\Delta G_i^{ab} + \Delta\Delta G_{A,i}^{ab} - \langle \Delta G_{j,k}^{Ab} \rangle$ |
| $aB$ | $\alpha_{ab} + \beta_B$ | $\Delta G_i^{ab} + \Delta\Delta G_{B,i}^{ab} - \langle \Delta G_{j,k}^{aB} \rangle$ |
| $AB$ | $\alpha_{ab} + \beta_A + \beta_B + \beta_{AB}$ | $\Delta G_i^{ab} + \Delta\Delta G_{A,i}^{ab} + \Delta\Delta G_{B,i}^{ab} - \langle \Delta G_{j,k}^{AB} \rangle$ |

The first model term, *α*_*ab*_, describes the wildtype function:

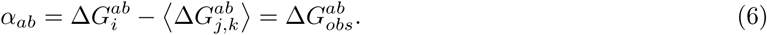

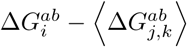 is the free energy difference between structure *i* and the Boltzmann-weighted ensemble of *j* and *k* (Equation 2). Put another way, *α*_*ab*_ is the measured Δ*G*_*obs*_ of the wildtype genotype.

Substituting the value of *α*_*ab*_ from Equation 6 into the linear models for genotypes *Ab* and *aB* in Table 1 and simplifying leads to mathematical descriptions of *β*_*A*_ and *β*_*B*_:

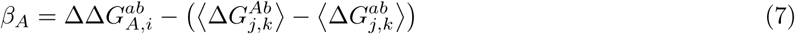

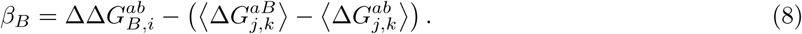

*β*_*A*_ and *β*_*B*_ describe the change in Δ*G*_*obs*_ upon introducing mutations. This is determined by the effect of the mutation on structure *i*, less the effect on *j* and *k*. We cannot directly estimate the effects of mutations on structure *i* because 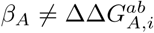. Rather, we only have access to the *difference* in the effect of a mutation between *i* and the other structures.

Finally, we determined the meaning of *β*_*AB*_ by substituting our values for *α*_*ab*_, *β*_*A*_, *β*_*B*_ (Equations 6, 7, and 8) and rearranging:

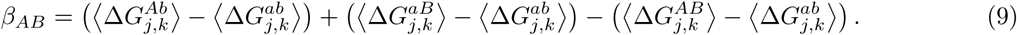

*β*_*AB*_ measures the change in free energy of *j* and *k* when mutations are introduced individually versus together. It is determined by the effects of mutations on structures *j* and *k*, not our observable structure *i*. This is because perturbations to the relative populations of *j* and *k* lead to nonlinear changes in the population of *i*, even if the mutations have no direct effect on the *i* structure. We find a similar term, *β*_*ABC*_, for three-way interaction between mutations, indicating that the thermodynamic ensemble can lead to high-order epistasis in addition to pairwise epistasis (see supplemental text).

### Requirements for ensemble epistasis

We next asked under what conditions ensemble epistasis is expected to arise. *β*_*AB*_ depends on the relative stabilities of *j* and *k* (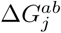 and 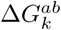) and the effects of mutations on *j* and *k* (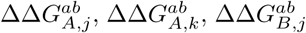, and 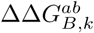) (Equations 5, 9). We found analytically (see supplemental text) that ensemble epistasis requires two conditions be met:

1. The protein populates at least two structures besides the observable structure *i*.
2. Mutations have differential effects on structures *j* and *k*.

To investigate what these conditions mean in practice, we calculated ensemble epistasis as a function of the relative populations of structures *j* and *k* 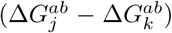 and the differential effects of mutations on structures *j* and *k* 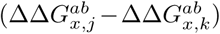 (Fig 2A). In panels B-C, we reveal the underlying ensemble that leads to the epistasis observed in Fig 2A. Notably, the length of the pink arrows illustrates the effect of mutation A in each genetic background, *ab* and *aB*. A difference in the length of the length of the pink arrows for the *ab* → *Ab* and *aB* → *AB* genotypes indicates epistasis.

**Fig 2:**
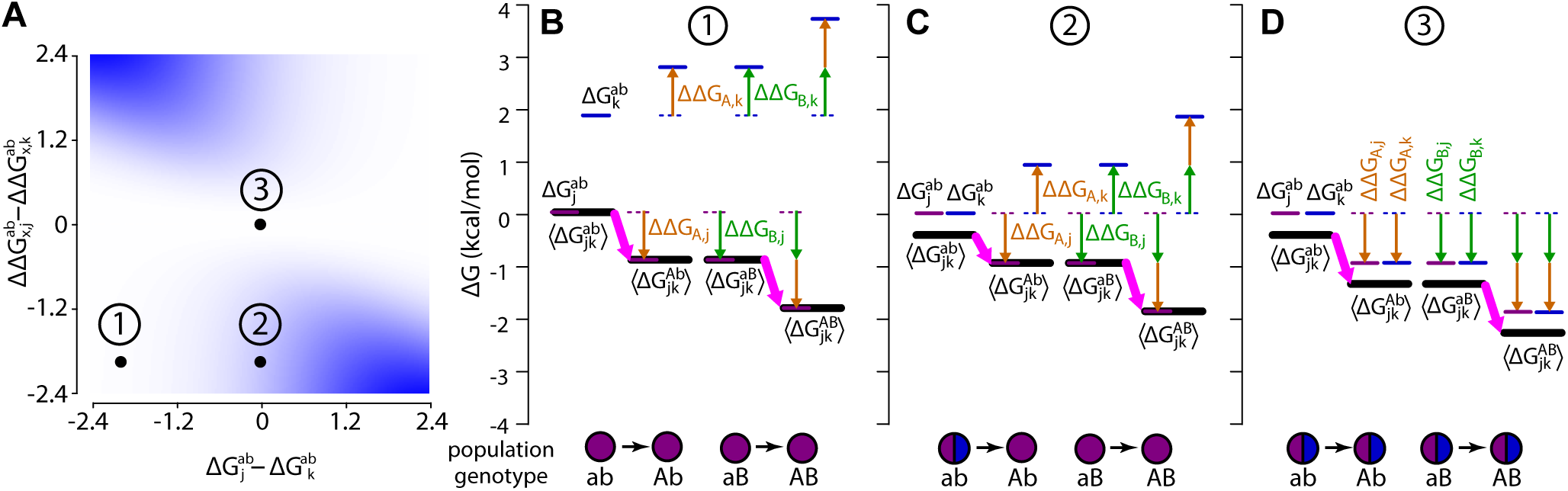
Ensemble epistasis arises from redistributed structural probabilities. A) Differential effects of the mutations *A* and *B* on structures *j* and *k* (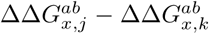 in *kcal* · *mol*^−1^, y-axis) as a function of the difference in the stability of structures *j* and *k* for the *ab* genotype (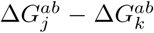 in *kcal* · *mol*^−1^, x-axis). Color indicates the magnitude of epistasis, ranging from 0 (white) to 1.6 *kcal* · *mol*^−1^ (blue). For the whole plot, *A* and *B* had identical effects (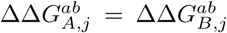 and 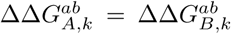). We set 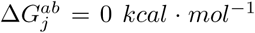 and 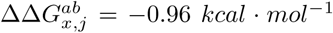 and then varied 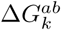 and 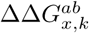 to sample parameter space. Panels B-C show the thermodynamic origins for the observed epistasis at points #1 (B), #2 (C), and #3 (D) indicated on panel A. The color scheme is consistent throughout: purple and blue lines are the energies of structures *j* and *k*, respectively; orange arrows show the effects of mutation *A*; green arrows show the effects of mutation *B*; heavy black lines are the observable ⟨Δ*G*_*jk*_⟩; heavy pink arrows are the observed effect of mutation *A* in the genotype indicated below the plot. The relative population of structures *j* and *k* are shown as a pie chart below the energy diagram.

We can see why multiple structures are required for ensemble epistasis by comparing points #1 and #2 on Fig 2A. At point #1, only structure *j* is populated for all genotypes (pie charts, Fig 2B); at point #2, structures *j* and *k* have equal starting populations (pie charts, Fig 2C). This difference in the starting populations of *j* and *k* leads to different epistatic outcomes. At point #1, both *ab* → *Ab* and *aB* → *AB* depend only on the effect of mutation *A* on structure *j* because it is the only structure populated. The lengths of the pink arrows are equal, indicating that there is no epistasis. At point #2, the effect of *ab* → *Ab* on ⟨Δ*G*_*j,k*_⟩ is moderate because the stabilization of structure *j* is offset by the entropic cost of depopulating structure *k*. When *A* is introduced second, *B* has already depopulated structure *k*. As a result, the effect of *aB* → *AB* is determined solely by its stabilization of structure *j*, and is thus larger than *ab* → *Ab*.

We can see why differential effects for each mutation are required by comparing points #2 and #3 on Fig 2A. At both points, structures *j* and *k* have equal starting populations (pie charts, Fig 2 C-D). At point #2, the mutations have opposite effects on structures *j* and *k* (Fig 2C); at point #3, the mutations have identical effects on structures *j* and *k* (Fig 2D). This means that for point #3 the introduction of *A* or *B* shifts the total energy landscape, but does not change the relative proportions of *j* and *k*. As a result, *A* has the same effect regardless of background (compare pink arrows, Fig 2D).

### Ensembles can lead to magnitude epistasis, sign-epistasis, reciprocal sign-epistasis

We next asked if the ensemble could lead to different evolutionarily-relevent classes of epistasis: magnitude, sign, and reciprocal sign. In magnitude epistasis, only the magnitude of a mutation’s effect changes when another mutation is introduced. In sign epistasis, the same mutation has a positive effect in one background and a negative effect in another. Finally, in reciprocal sign epistasis, both mutations exhibit sign epistasis.

In Fig 3A, we set the initial energies of structures *j* and *k* to be equal 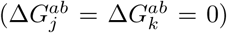. We then slid along values for the difference in the effects of mutations *A* and *B* on *j* and *k*. We found four regimes, corresponding to magnitude, reciprocal sign, sign, and no epistasis. The effect of mutation *A* on each structure is shown in Fig 3B; it destabilizes structure *j* by 0.35 *kcal* · *mol*^−1^ while stabilizing *k* by the same amount, leading to a decrease in the observable, 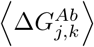. We held the difference in the effect of mutation *A* on *j* and *k* constant at 0.7 *kcal* · *mol*^−1^.

**Fig 3:**
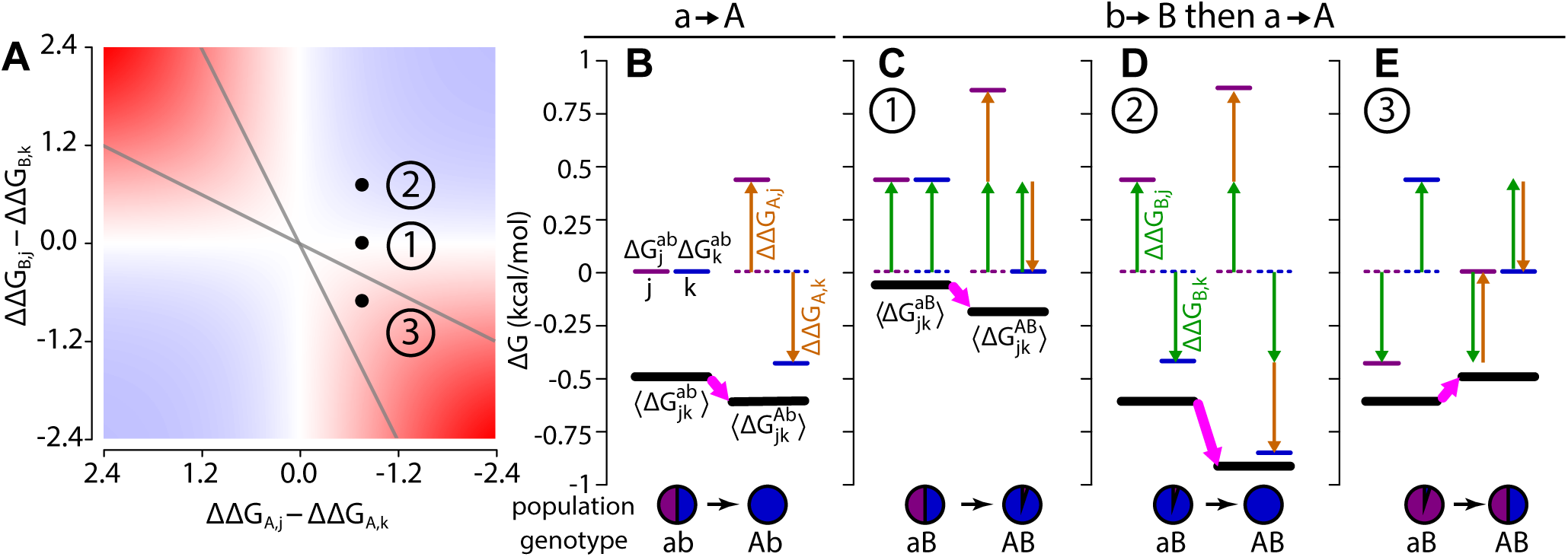
Ensemble epistasis arises when mutations have different effects on different structures. A) Epistasis calculated for a three-structure ensemble that starts with 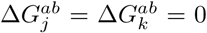. The differences in the effects of mutations *A* and *B* on structures *j* and *k* are indicated on the *x*- and *y*-axes. The magnitude of epistasis is indicated by the color, ranging from − 1.6 (dark red) to 0 (white) to +1.6 *kcal* · *mol*^−1^ (dark blue). Grey lines delineate regions of reciprocal sign (red regions within the lines) and sign epistasis (red regions outside of the lines). Panels B-E show the thermodynamic origins of the epistasis indicated by points 1, 2 and 3 on panel A. The effect of mutation *A* is constant in all panels; the effect of mutation *B* differs depending on the scenario. The color scheme is consistent with Fig 2. B) The effect of *A* introduced into the *ab* background. *A* destabilizes *j* and stabilizes *k*, stabilizing the observable. C) Scenario 1: *A* and *B* exhibit no epistasis. Mutation *B* has the same effect on structures *j* and *k*. D) Scenario 2: *A* and *B* act synergistically to destabilize *j* and stabilize *k*. E) Scenario 3: Mutations *A* and *B* have opposite effects on structures *j* and *k*.

If *A* or *B* have the same effect on structures *j* and *k*, we see no epistasis (the white strips, Fig 3A). In Fig 3C, mutation *B* destabilizes both *j* and *k* by 0.35 *kcal*· *mol*^−1^. Because mutation *B* does not have differential effects on each structure, ⟨Δ*G*_*j,k*_⟩ is globally shifted by +0.35 *kcal* · *mol*^−1^. Introducing *A* and *B* together yields no epistasis because both the *ab* and *aB* genotypes have identical configurations—the observed effect comes only from mutation *A* (compare pink arrows in Fig 3B and Fig 3C).

When mutations are both stabilizing or both destabilizing, we observe magnitude epistasis (blue regions, Fig 3A). In Fig 3D, mutations *A* and *B* have synergistic effects on each structure: *k* is stabilized while *j* is destabilized. We see magnitude epistasis because although the relative population of *j* is reduced, it still has weight in the Boltzmann-weighted average stability (compare pink arrows in Fig 3B and 3D).

When both mutations have opposite effects, we observe reciprocal sign and sign epistasis (red regions, Fig 3A). Fig 3E shows an example of reciprocal sign epistasis. Mutations *A* and *B* have individually stabilizing effects on the observable but are destabilizing when combined. This occurs because *A* and *B* have opposite effects on *j* and *k*—*A* destabilizes *j* and stabilizes *k*, while *B* stabilizes *j* and destabilizes *k*. The effects are equal in magnitude but opposite in sign so their combined effects cancel, yielding an observable equal to that of the *ab* genotype (compare pink arrows in Fig 3B and 3E).

### Ensemble epistasis may be a common feature in protein mutant cycles

Above we showed mathematically that ensemble epistasis can arise when two conditions are met. Is it plausible that these conditions are met by real macromolecules, where the the effects of mutations may be corrleated between structures? What is the magnitude of the resulting epistasis? How might one detect this effect experimentally?

We investigated these questions using the allosteric *Ca*^2+^ signaling protein, human S100A4. The protein adopts a three-structure ensemble (Fig 4A) [19–21]. In the absence of *Ca*^2+^, it favors the “*apo*” structure (Fig 4A, slate); addition of *Ca*^2+^ stabilizes the “*ca*” structure with an exposed hydrophobic peptide-binding surface (Fig 4A, purple); finally, addition of *peptide* leads to formation the “*capep*” structure that has both *Ca*^2+^ and *peptide* bound (Fig 4A, green).

**Fig 4:**
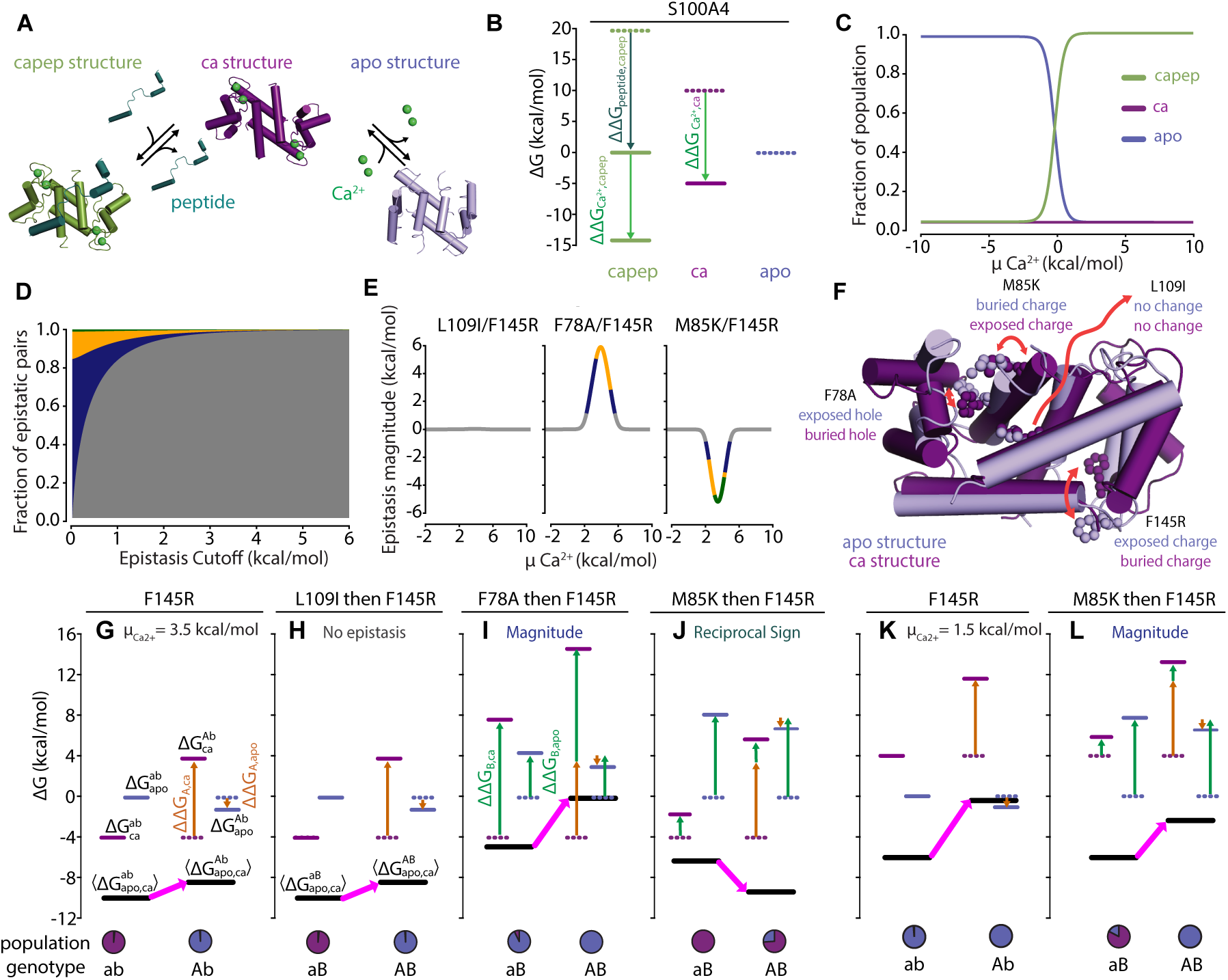
The ensemble of S100A4 exhibits ensemble epistasis. A) Three-structure ensemble of the S100A4 protein. The apo structure (*apo*, PDB: 1M31, slate) is in equilibrium with the *Ca*^2+^ bound (*ca*, PDB: 2Q91, purple) and *Ca*^2+^/*peptide* bound (*capep*, PDB: 5LPU, green) structures when *Ca*^2+^ (lime green spheres) and *peptide* (teal) are present. B) Assigned energies (*kcal* · *mol*^−1^) of S100A4 structures. *Apo* is most stable when *peptide, µ*_*peptide*_, and *Ca*^2+^ chemical potentials, 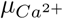, are zero (dashed lines). *Capep* is stabilized by increasing *µ*_*peptide*_ = 20 *kcal* · *mol*^−1^ (teal arrow, solid green line). Increasing 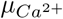 alters the energies of both *ca* and *capep* (lime green arrow, solid lines). C) Relative population (y-axis) of each structure as a function of 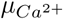 (*kcal* · *mol*^−1^, x-axis) at *µ*_*peptide*_ = 20 *kcal* · *mol*^−1^. D) Fractional contribution of each epistatic type (y-axis) as a function of epistatic magnitude cutoff (*kcal* · *mol*^−1^, x-axis), colored by type: reciprocal sign (teal), sign (gold), and magnitude (dark blue). Pairs with maximum epistasis below the cutoff compose the no epistasis fraction (grey). E) Epistatic magnitude (*kcal* · *mol*^−1^, y-axis) as a function of 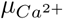 (*kcal* · *mol*^−1^, x-axis) for three mutation pairs: L109I/F145R (left panel), F78A/F145R (middle panel), and M85K/F145R/M85K (right panel). Color is consistent with epistatic type from panel D. F) Positions of mutations in the *apo* (slate) and *ca* (purple) structures. Text indicates their relative environments in each structure. Red arrows indicate changes in position. G-L) Thermodynamic origins of epistasis for three mutation pairs at 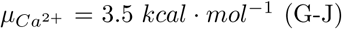 or 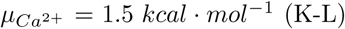. Mutation *A* (F145R) is constant; mutation *B* differs in panels H-J and L. The color scheme is consistent throughout: blue and purple lines are the energies of *apo* and *ca*, respectively; all other colors are consistent with Fig 2-3. Specific mutations and epistatic classes are indicated at the top of the panel; genotypes and relative populations are below. G) Introduction of mutation *A* (F145R) into the wildtype *ab* background at 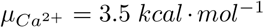. H) No epistasis scenario: mutations F145R (*a* → *A*) and L109I (*b* → *B*). I) Magnitude epistasis scenario: mutations F145R (*a* → *A*) and F78A (*b* → *B*). I) Reciprocal sign epistasis scenario: mutations F145R (*a* → *A*) and M85K (*b* → *B*). K) Introduction of mutation *A* (F145R) into the *ab* background at 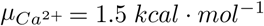) Magnitude epistasis scenario: mutations F145R (*a* → *A*) and M85K (*b* → *B*).

To model the ensemble, we selected reference concentrations of *Ca*^2+^ and *peptide* such that 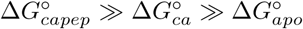 (Fig 4B). We know experimentally that the protein favors the *apo* conformation in the absence of *Ca*^2+^ and *peptide*. We then modeled the signaling behavior of S100A4 by changing the concentrations of *Ca*^2+^ and *peptide*: 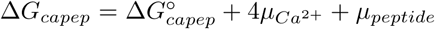 and 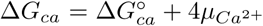, where 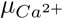 and *µ*_*peptide*_ are the chemical potentials of *Ca*^2+^ and *peptide* relative to their reference concentrations (Fig 4B). Depending on our choice of 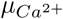 and *µ*_*peptide*_, we can observe different relative populations of the *apo, ca*, and *capep* structures (Fig 4C). We treated the *capep* structure as our “active” structure, labelled “*i*” in Fig 1D and Equation 2. Ensemble epistasis depends solely on unobserved states (Equation 9). Thus, we focused our remaining analysis only on the *apo* and *ca* structures.

We next used ROSETTA to estimate the stability effects of all possible single point mutations to the *apo* and *ca* structures of S100A4. This gives us ΔΔ*G*_*x,apo*_ and ΔΔ*G*_*x,ca*_ for every mutation. We then calculated epistasis in Δ*G*_*obs*_, as a function of 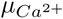 and *µ*_*peptide*_, for all 5.6 million pairs of these mutations using Equation 9. See methods for details.

We next analyzed the magnitude of ensemble epistasis throughout our dataset. We considered each pair’s maximum value of epistasis within our 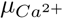 range. Roughly 27% of pairs generated epistasis on the order of thermal fluctuation, 0.6 *kcal* · *mol*^−1^ (Fig 4D). We found that 18% of pairs exhibited magnitude-, 8% sign-, and 1% reciprocal-sign-epistasis at this cutoff. About 5% of pairs exhibited epistasis with a magnitude above 2 *kcal* · *mol*^−1^.

We observed three basic patterns of 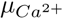-dependent epistatic magnitude, as exemplified by the three mutant pairs shown in Fig 4E: F145R/L109I had no epistasis (left panel) while F145R/F78A had positive epistasis (middle panel) and F145R/M85K had negative epistasis (right panel). We observed peaks in epistasis at intermediate values of 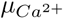, where the *apo, ca*, and *capep* structures may all be populated. In contrast, we observed no epistasis at low 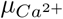 (where only the *apo* conformation is populated) or high 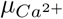 (where only the *capep* structure is populated). Interestingly, the type of epistasis observed—magnitude (dark blue), sign (gold), or reciprocal sign (teal)—was also dependent upon 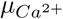 (Fig 4E). This was quite common in our dataset: approximately 68% of pairs with an epistatic magnitude above 0.6 *kcal·mol*^−1^ switched epistatic type at least once as 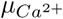 increased.

To understand the structural origins of the observed epistasis, we compared the the positions of each mutation from Fig 4E in the *apo* (slate, Fig 4F) and *ca* (purple, Fig 4F) structures. First we consider F145R. F145R is solvent exposed in the *apo* structure but buried in the *ca* structure. As a consequence, introducing Arg mildly stabilizes the *apo* structure, but dramatically destabilizes the *ca* structure due to burying its charge. L109I is a conservative mutation at a site whose environment is essentially unchanged between the *apo* and *ca* structure. F78A is solvent exposed in the *apo* structure, but buried in the *ca* structure. The Phe to Ala mutation is destabilizing to the *ca* structure due to the loss of hydrophobic contacts. Finally, M85K is buried in the *apo* structure, but exposed in the *ca* structure. Mutation to Lys introduces a buried charge, greatly destabilizing it due to the cost of ion desolvation.

The differences in the effect of L109I, F78A, and M85K cause them to exhibit different types of epistasis when paired with F145R. F145R exhibits no epistasis when paired with L109I at 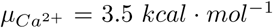 (Fig 4H). The L109I mutation has a negligible effect on the *apo* and *ca* structures (genotype *aB*, Fig 4H). As a result, F145R has the same effect on ⟨Δ*G*_*apo,ca*_⟩ when it is introduced into L109I and wildtype backgrounds (compare pink arrows in Fig 4G and 4H).

Pairing F145R with F78A results in magnitude epistasis. F78A is destabilizing to both structures, but much more so to the *ca* structure (genotype *aB*, Fig 4I). Both F78A and F145R preferentially destabilize the *ca* structure, leading to a dramatic decrease in its relative population when introduced together (green arrows, Fig 4I). Synergistic destabilization of the *ca* structure and the resultant increase in ⟨Δ*G*_*apo,ca*_⟩ leads to strong magnitude epistasis (compare pink arrows in Fig 4G and 4I).

F145R exhibits reciprocal sign epistasis when paired with M85K. The Lys mutation is greatly destabilizing to the *apo* structure while it has a much smaller destabilizing effect on the *ca* structure (green arrows, Fig 4J). The net effect of M85K is a decrease in the stability of ⟨Δ*G*_*apo,ca*_⟩. Combining both individually destabilizing mutations restores the relative energies of the excited structures to a globally shifted wildtype configuration, leading to a net stabilization of ⟨Δ*G*_*apo,ca*_⟩ and the observation of reciprocal sign epistasis (pink arrows, Fig 4J).

Intriguingly, a slight decrease from 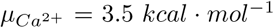 in panels G-J to 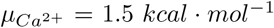 switches the type of epistasis from reciprocal sign to magnitude for the F145R/M85K pair (Fig 4E and Fig 4K-L). The switch from reciprocal sign to magnitude epistasis is solely due to the change in the relative energies of the *apo* and *ca* structure in the *ab* genotype: although the wildtype configuration is restored, the *ca* structure is so severely destabilized that it has a negligible impact on the observable (compare pink arrows in Fig 4K and Fig 4L).

Effector- or environment-dependent epistasis may signal ensemble epistasis. We propose that one might experimentally test for it as we did above: by altering the relative populations of the excited structures by varying effector concentration. Ensemble epistasis should be maximized at concentrations where many distinct structures are populated (i.e. at concentrations where functional transitions occur) and minimized when mutations can impact only a single state (i.e. low 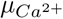).

## Discussion

### Ensemble epistasis should be pervasive in biology

We find that epistasis can arise from a fundamental property of macromolecules: the thermodynamic ensemble. Previously we observed ensemble epistasis using lattice models, but the conditions under which it arises and if they are plausibly met outside of toy systems remained unresolved [14]. Here we used a simple—but general—thermodynamic model to study the nonlinear mapping between mutational effects and an observable. Nonlinearity arises because mutations can affect any structure in the ensemble. We observe epistasis because observables are averaged over the entire ensemble and cannot be separated into additive components.

We expect ensemble epistasis in systems where 1) at least three structures are populated and 2) mutations have differential effects on excited structures (Fig 2). The first requirement is likely common; multi-structure ensembles often underlie biological function—from allostery to fold-switching (Fig 1D) [16]. To address the plausibility of these requirements in a real protein, we modeled the effects of mutations on the three-structure ensemble of the allosteric *Ca*^2+^ signalling protein S100A4. This meets the first requirement. We then identified mutations that met the second requirement and found that they resulted in epistasis, suggesting that—at least in principle—ensemble epistasis should be detectable in real proteins (Fig 4D).

There is mounting indirect evidence of links between epistasis and thermodynamic ensembles, suggesting the second requirement is plausible in macromolecules. For example, in TEM-1 Beta-lactamase, two adaptive mutations were identified that independently increased structural heterogeneity and function. Together the mutations exhibited epistasis, shifting the ensemble into a dominantly non-productive structure [22]. Epistasis also underlies changes in dynamics that caused divergence between Src and Abl kinases and the evolution of fold-switching proteins [23, 24].

### Ensemble epistasis may shape evolution

We have shown that simple ensembles give rise to magnitude, sign, reciprocal sign, and high-order epistasis (Fig 3, supplemental text). Sign and reciprocal sign epistasis are particularly important; they can decrease accessible evolutionary trajectories and are required for the presence of multiple peaks in fitness landscapes [9, 10, 25–32]. High-order epistasis can alter accessibility and can facilitate bypassing evolutionary dead-ends in genotype-phenotype maps, making evolution deeply unpredictable [7, 8, 33, 34].

We saw 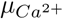-dependent epistasis in S100A4 with a peak at regions where multiple structures were populated. 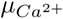-dependent changes in epistatic magnitude were often accompanied by switches in the observed type of epistasis. A similar phenomenon has been observed in allosteric proteins where ligands can act as agonists or antagonists in response to changes in environment, ultimately via changes in the thermodynamic ensemble [35]. Intriguingly, reshaping the thermodynamic ensemble has the largest epistatic effect—both in magnitude and type—under physiologically relevant conditions, precisely where macromolecules toggle between distinct modes of function. Ensemble epistasis may represent an evolutionary mechanism that serves to couple environmental changes to increases in phenotypic variation, possibly increasing evolvability.

Studies have linked the thermodynamic ensemble to evolvability, but the role of epistasis is often unclear [36, 37]. A recent study showed that epistasis between two dynamics-altering mutations in TEM-1 Beta-lactamase facilitated the evolution of distinct enzymatic activities [38]. There are also examples of ensemble-facilitated evolution requiring minimal epistasis [36]. The reduced role of ensemble epistasis may be related to stability-mediated thresholds, where mutations are additive until destabilizing mutations accumulate such that the threshold required for function is crossed [36, 39, 40]. Notably, the mathematical framework of the thermodynamic ensemble is not limited to proteins—it has been used to describe much more complex biological systems like signalling networks and bacterial communities [41–46].

### Detecting ensemble epistasis

Here we assumed a discretely defined, three-structure system: one structure (*i*) that determines the biological observable of interest and two excited structures (*j* and *k*) that give rise to ensemble epistasis. While this assumption will be unlikely to hold for many biological systems, our mathematical formalism above can be extended to *n* observable structures and *m* excited structures and still give rise to ensemble epistasis when the two minimal conditions are met.

A straightforward experimental test for ensemble epistasis would be to perturb the thermodynamic ensemble by tuning environmental parameters. We observed effector-dependent epistasis in the S100A4 protein, where epistasis changed with the addition of *Ca*^2+^. The magnitude of epistasis peaked in regions where mutations were able to affect multiple structures, or where the chances of satisfying the requirements are the highest. Environmental-dependent epistasis has been noted previously, pointing to a possible underlying ensemble structure [32, 47–53].

### Modeling ensemble epistasis

Some important goals of in modeling epistasis are to understand evolution, interpret high-throughput experiments, and engineer macromolecules. We see two general strategies to model ensemble epistasis in these contexts: 1) account for it with specific mechanistic models and 2) embrace the statistical nature of ensembles and model epistasis as a Gaussian noise term [54]. Recently, the first approach was taken to decompose mutational effects in the GB1 protein [55]. A three-structure ensemble model was able to explain much of the epistasis observed in the dataset. The remaining epistasis pointed towards residues that contribute to functionally important structural dynamics. This approach both reduces nonlinearity and gives mechanistic information about the system.

The above approach may work for simple, well-behaved proteins. For complex proteins, ensemble epistasis might be better modeled as a Gaussian noise term. Attempts to model a fundamentally statistical, population-level phenomenon with specific epistatic coefficients may be misleading. This has been noted previously but has foundations in the mathematics of thermodynamic ensembles: the Boltzmann-distribution collapses to a Gaussian when many structures are populated [54]. The statistical nature of thermodynamic ensembles likely means they cannot be distilled into a predictive mechanistic framework.

## Conclusion

While the ubiquity of ensemble epistasis remains unknown, we anticipate that it is widespread. Thermodynamic ensembles—a fundamental property of macromolecules—can give rise to epistasis of all types, namely magnitude, sign, and reciprocal sign, suggesting that it can shape how macromolecules evolve. Collectively, this points to ensemble epistasis as a potentially universal mechanism of epistasis, making contingency and unpredictability a fixed feature of biology.

## Methods

All analyses and ROSETTA input files can be downloaded from https://github.com/harmslab/ensemble_epistasis.

For the S100A4 epistasis analysis, we used three published structures for S100A4: the apo structure (PDB 1M31), the *Ca*^2+^ bound structure (PDB 2Q91), and the structure bound to both *Ca*^2+^ and a peptide extracted from Annexin A2 (PDB 5LPU). We removed all non-*Ca*^2+^ small molecules (including waters) and edited the files to have an identical set of non-hydrogen atoms for the S100A4 chains (trimming any residues before alanine 2 and after phenylalanine 93 in the uniprot sequence, P26447). We arbitrarily selected the first NMR model for the apo structure. Using ROSETTA (Linux build 2018.33.60351), we generated five independent, pre-minimized structures for each of the states (*apo, Ca*^2+^, and *Ca*^2+^/*peptide*). We then used the “cartesian_ddg” binary to introduce each mutation three times into each of these five pre-minimized structures, yielding 15 calculated ΔΔ*G* values for each mutation in each of the three states [56]. Finally, we averaged the 15 values for each mutation in each state. We assumed the units of these ΔΔ*G* values were in *kcal* · *mol*^−1^ [57].

For a given genotype, we described the free energy of the calcium-bound form as a function of calcium chemical potential 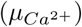 with the expression 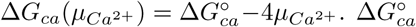 is a constant describing both the relative stability of the “open” form of the protein relative to the “closed” form and the affinity of the open form for *Ca*^2+^. We treated the free energy of the *apo* form as 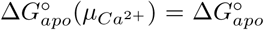, where 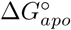 measures the free energy of the *apo* form relative to an arbitrary reference state. For convenience, we set 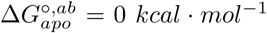 and 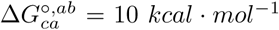 for *µ* = 0 *kcal* · *mol*^−1^. This models the fact that, at some reference [*Ca*^2+^], the closed form is favored over “open” form. As [*Ca*^2+^] increases, 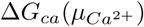 becomes more negative and eventually becomes more favorable tham Δ*G*_*apo*_.

We modeled the effects of mutations as changes to 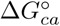 and 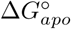. For the *Ab* genotype, for example, we would write:

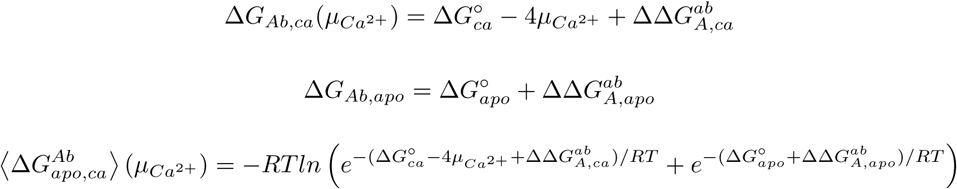

were ΔΔ*G*_*A,ca*_ and ΔΔ*G*_*A,apo*_ are the energetic effects of mutation *A* and the calcium and apo states, respectively. See the supplemental text for further information, including a derivation of the model.

