## Supplemental Text for "Ensemble epistasis: thermodynamic origins of non-additivity between mutations"

### 1 Ensemble epistasis requires three or more structures:

The definition of the Boltzmann-weighted stability of structures $j$ and $k$ for some genotype is (Equation 3):

$$\left\langle \Delta G_{j,k}^{genotype} \right\rangle = -RTln\left(e^{-\Delta G_j^{genotype}/RT} + e^{-\Delta G_k^{genotype}/RT}\right)$$

If $\Delta G_k \gg \Delta G_j$, structure $k$ is not appreciably populated; we set the second term to 0.

$$\left\langle \Delta G_{j,k}^{genotype} \right\rangle = -RTln\left(e^{-\Delta G_j^{genotype}/RT}\right)$$

$$\left\langle \Delta G_{j,k}^{genotype} \right\rangle = \Delta G_j^{genotype}$$

Now we have a two-state system, populating only structures $i$ and $j$. $\Delta G_j^{Ab}$ is given by the free energy of the wildtype structure perturbed by mutation $A$:

$$\Delta G_j^{Ab} = \Delta G_j^{ab} + \Delta\Delta G_{A,j}^{ab}.$$

Substituting expressions for the free energy of each genotype into our expression for $\beta_{AB}$, we find that all of the terms cancel:

$$\beta_{AB} = \left(\Delta G_j^{ab} + \Delta\Delta G_{A,j}^{ab} - \Delta G_j^{ab}\right) + \left(\Delta G_j^{ab} + \Delta\Delta G_{B,j}^{ab} - \Delta G_j^{ab}\right) - \left(\Delta G_j^{ab} + \Delta\Delta G_{A,j}^{ab} + \Delta\Delta G_{B,j}^{ab} - \Delta G_j^{ab}\right),$$

yielding

$$\beta_{AB} = 0.$$

Thus, two-state systems cannot exhibit ensemble epistasis.

### 2 Ensemble epistasis requires mutations have differential effects:

We set $\Delta\Delta G_{B,j}^{ab} = \Delta\Delta G_{B,k}^{ab} = \Delta\Delta G_B^{ab}$ for mutation $B$, while allowing mutation $A$ to have different effects on each structure: $\Delta\Delta G_{A,j}^{ab} \neq \Delta\Delta G_{A,k}^{ab}$.

Consider the effect of $B$ in the $ab$ background:

$$\left\langle \Delta G_{j,k}^{aB} \right\rangle = -RTln\left(e^{-\left(\Delta G_j^{ab} + \Delta\Delta G_B^{ab}\right)/RT} + e^{-\left(\Delta G_k^{ab} + \Delta\Delta G_B^{ab}\right)/RT}\right)$$

We can factor out $\Delta\Delta G_{B}^{ab}$:

$$\left\langle \Delta G_{j,k}^{aB} \right\rangle = -RTln\left(e^{-\Delta\Delta G_{B}^{ab}/RT}e^{-\Delta G_{j}^{ab}/RT} + e^{-\Delta\Delta G_{B}^{ab}/RT}e^{-\Delta G_{k}^{ab}/RT}\right)$$

$$\left\langle \Delta G_{j,k}^{aB} \right\rangle = -RTln\left(e^{-\Delta\Delta G_{B}^{ab}/RT}\left(e^{-\Delta G_{j}^{ab}/RT} + e^{-\Delta G_{k}^{ab}/RT}\right)\right)$$

$$\left\langle \Delta G_{j,k}^{aB} \right\rangle = -RTln\left(e^{-\Delta\Delta G_{B}^{ab}/RT}\right) - RTln\left(e^{-\Delta G_{j}^{ab}/RT} + e^{-\Delta G_{k}^{ab}/RT}\right)$$

$$\left\langle \Delta G_{j,k}^{aB} \right\rangle = \Delta\Delta G_{B}^{ab} - RTln\left(e^{-\Delta G_{j}^{ab}/RT} + e^{-\Delta G_{k}^{ab}/RT}\right)$$

$$\left\langle \Delta G_{j,k}^{aB} \right\rangle = \Delta\Delta G_{B}^{ab} + \left\langle \Delta G_{j,k}^{ab} \right\rangle.$$

Consider the effect of $B$ in the $Ab$ background:

$$\left\langle \Delta G_{j,k}^{AB} \right\rangle = -RTln\left(e^{-\left(\Delta G_{j}^{ab}+\Delta\Delta G_{A,j}^{ab}+\Delta\Delta G_{B}^{ab}\right)/RT} + e^{-\left(\Delta G_{k}^{ab}+\Delta\Delta G_{A,k}^{ab}+\Delta\Delta G_{B}^{ab}\right)/RT}\right).$$

Again, we factor out $\Delta\Delta G_{B}^{ab}$:

$$\left\langle \Delta G_{j,k}^{AB} \right\rangle = -RTln\left(\left(e^{-\Delta\Delta G_{B,j}^{ab}/RT}\right)e^{-\left(\Delta G_{j}^{ab}+\Delta\Delta G_{A,j}^{ab}\right)/RT} + \left(e^{-\Delta\Delta G_{B,j}^{ab}/RT}\right)e^{-\left(\Delta G_{k}^{ab}+\Delta\Delta G_{A,k}^{ab}\right)/RT}\right)$$

$$\left\langle \Delta G_{j,k}^{AB} \right\rangle = -RTln\left(\left(e^{-\Delta\Delta G_{B,j}^{ab}/RT}\right)\left(e^{-\left(\Delta G_{j}^{ab}+\Delta\Delta G_{A,j}^{ab}\right)/RT} + e^{-\left(\Delta G_{k}^{ab}+\Delta\Delta G_{A,k}^{ab}\right)/RT}\right)\right)$$

$$\left\langle \Delta G_{j,k}^{AB} \right\rangle = -RTln\left(e^{-\Delta\Delta G_{B,j}^{ab}/RT}\right) - RTln\left(e^{-\left(\Delta G_{j}^{ab}+\Delta\Delta G_{A,j}^{ab}\right)/RT} + e^{-\left(\Delta G_{k}^{ab}+\Delta\Delta G_{A,k}^{ab}\right)/RT}\right)$$

$$\left\langle \Delta G_{j,k}^{AB} \right\rangle = \Delta\Delta G_{B,j}^{ab} - RTln\left(e^{-\left(\Delta G_{j}^{ab}+\Delta\Delta G_{A,j}^{ab}\right)/RT} + e^{-\left(\Delta G_{k}^{ab}+\Delta\Delta G_{A,k}^{ab}\right)/RT}\right)$$

$$\left\langle \Delta G_{j,k}^{AB} \right\rangle = \Delta\Delta G_{B,j}^{ab} + \left\langle \Delta G_{j,k}^{Ab} \right\rangle.$$

Substitute expressions for $\left\langle \Delta G_{j,k}^{aB} \right\rangle$ and $\left\langle \Delta G_{j,k}^{AB} \right\rangle$ into the expression for $\beta_{AB}$ (Equation 9):

$$\beta_{AB} = \left(\left\langle \Delta G_{j,k}^{Ab} \right\rangle - \left\langle \Delta G_{j,k}^{ab} \right\rangle\right) + \left(\left\langle \Delta G_{j,k}^{aB} \right\rangle - \left\langle \Delta G_{j,k}^{ab} \right\rangle\right) - \left(\left\langle \Delta G_{j,k}^{AB} \right\rangle - \left\langle \Delta G_{j,k}^{ab} \right\rangle\right)$$

$$\beta_{AB} = \left(\left\langle \Delta G_{j,k}^{Ab} \right\rangle - \left\langle \Delta G_{j,k}^{ab} \right\rangle\right) + \left(\Delta\Delta G_{B}^{ab} + \left\langle \Delta G_{j,k}^{ab} \right\rangle - \left\langle \Delta G_{j,k}^{ab} \right\rangle\right) - \left(\Delta\Delta G_{B,j}^{ab} + \left\langle \Delta G_{j,k}^{Ab} \right\rangle - \left\langle \Delta G_{j,k}^{ab} \right\rangle\right).$$

All terms cancel, yielding:

$$\beta_{AB} = 0.$$

Mutations must have differential effects on two ore more structures to observe ensemble epistasis.

### 3 Ensembles can lead to high-order epistasis

In this, we consider mutations at three sites as an extension of the two-site case. Below is a table describing the three-mutation case, directly paralleling the two-site case in Table 1.

| genotype | $\Delta G$ | model |
|---|---|---|
| $abc$ | $\alpha_{abc}$ | $\Delta G_i^{abc} - \left\langle \Delta G_{j,k}^{abc} \right\rangle$ |
| $Abc$ | $\alpha_{abc} + \beta_A$ | $\Delta G_i^{abc} + \Delta\Delta G_{A,i}^{abc} - \left\langle \Delta G_{j,k}^{Abc} \right\rangle$ |
| $aBc$ | $\alpha_{abc} + \beta_B$ | $\Delta G_i^{abc} + \Delta\Delta G_{B,i}^{abc} - \left\langle \Delta G_{j,k}^{aBc} \right\rangle$ |
| $abC$ | $\alpha_{abc} + \beta_C$ | $\Delta G_i^{abc} + \Delta\Delta G_{B,i}^{abc} - \left\langle \Delta G_{j,k}^{abC} \right\rangle$ |
| $ABc$ | $\alpha_{abc} + \beta_A + \beta_B + \beta_{AB}$ | $\Delta G_i^{abc} + \Delta\Delta G_{A,i}^{abc} + \Delta\Delta G_{B,i}^{abc} - \left\langle \Delta G_{j,k}^{ABc} \right\rangle$ |
| $AbC$ | $\alpha_{abc} + \beta_A + \beta_C + \beta_{AC}$ | $\Delta G_i^{abc} + \Delta\Delta G_{A,i}^{abc} + \Delta\Delta G_{C,i}^{abc} - \left\langle \Delta G_{j,k}^{AbC} \right\rangle$ |
| $aBC$ | $\alpha_{abc} + \beta_B + \beta_C + \beta_{BC}$ | $\Delta G_i^{abc} + \Delta\Delta G_{B,i}^{abc} + \Delta\Delta G_{C,i}^{abc} - \left\langle \Delta G_{j,k}^{aBC} \right\rangle$ |
| $ABC$ | $\alpha_{abc} + \beta_A + \beta_B + \beta_C + \beta_{AB} + \beta_{AC} + \beta_{BC} + \beta_{ABC}$ | $\Delta G_i^{abc} + \Delta\Delta G_{A,i}^{abc} + \Delta\Delta G_{B,i}^{abc} + \Delta\Delta G_{C,i}^{abc} - \left\langle \Delta G_{j,k}^{ABC} \right\rangle$ |

We can solve for each coefficient in the epistatic model in thermodynamic terms:

$$\alpha_{abc} = \Delta G_i^{abc} - \left\langle \Delta G_{j,k}^{abc} \right\rangle$$

$$\beta_A = \Delta\Delta G_{A,i}^{abc} - \left(\left\langle \Delta G_{j,k}^{Abc} \right\rangle - \left\langle \Delta G_{j,k}^{abc} \right\rangle\right)$$

$$\beta_B = \Delta\Delta G_{B,i}^{abc} - \left(\left\langle \Delta G_{j,k}^{aBc} \right\rangle - \left\langle \Delta G_{j,k}^{abc} \right\rangle\right)$$

$$\beta_C = \Delta\Delta G_{C,i}^{abc} - \left(\left\langle \Delta G_{j,k}^{abC} \right\rangle - \left\langle \Delta G_{j,k}^{abc} \right\rangle\right)$$

$$\beta_{AB} = \left(\left\langle \Delta G_{j,k}^{Abc} \right\rangle - \left\langle \Delta G_{j,k}^{abc} \right\rangle\right) + \left(\left\langle \Delta G_{j,k}^{aBc} \right\rangle - \left\langle \Delta G_{j,k}^{abc} \right\rangle\right) - \left(\left\langle \Delta G_{j,k}^{ABc} \right\rangle - \left\langle \Delta G_{j,k}^{abc} \right\rangle\right)$$

$$\beta_{AC} = \left(\left\langle \Delta G_{j,k}^{Abc} \right\rangle - \left\langle \Delta G_{j,k}^{abc} \right\rangle\right) + \left(\left\langle \Delta G_{j,k}^{abC} \right\rangle - \left\langle \Delta G_{j,k}^{abc} \right\rangle\right) - \left(\left\langle \Delta G_{j,k}^{AbC} \right\rangle - \left\langle \Delta G_{j,k}^{abc} \right\rangle\right)$$

$$\beta_{AC} = \left(\left\langle \Delta G_{j,k}^{aBc} \right\rangle - \left\langle \Delta G_{j,k}^{abc} \right\rangle\right) + \left(\left\langle \Delta G_{j,k}^{abC} \right\rangle - \left\langle \Delta G_{j,k}^{abc} \right\rangle\right) - \left(\left\langle \Delta G_{j,k}^{aBC} \right\rangle - \left\langle \Delta G_{j,k}^{abc} \right\rangle\right)$$

$$\begin{aligned}\beta_{ABC} = \; & -\left[\left(\left\langle \Delta G_{j,k}^{Abc} \right\rangle - \left\langle \Delta G_{j,k}^{abc} \right\rangle\right) + \left(\left\langle \Delta G_{j,k}^{aBc} \right\rangle - \left\langle \Delta G_{j,k}^{abc} \right\rangle\right) + \left(\left\langle \Delta G_{j,k}^{abC} \right\rangle - \left\langle \Delta G_{j,k}^{abc} \right\rangle\right)\right] + \backslash \\ & \left[\left(\left\langle \Delta G_{j,k}^{ABc} \right\rangle - \left\langle \Delta G_{j,k}^{abc} \right\rangle\right) + \left(\left\langle \Delta G_{j,k}^{AbC} \right\rangle - \left\langle \Delta G_{j,k}^{abc} \right\rangle\right) + \left(\left\langle \Delta G_{j,k}^{aBC} \right\rangle - \left\langle \Delta G_{j,k}^{abc} \right\rangle\right)\right] - \backslash \\ & \left[\left(\left\langle \Delta G_{j,k}^{ABC} \right\rangle - \left\langle \Delta G_{j,k}^{abc} \right\rangle\right)\right]\end{aligned}$$

This shows, directly analogous to the pairwise epistatic case, that nonlinear perturbations of mutations to unobserved structures $j$ and $k$ lead to a potentially non-zero three-way interaction term $\beta_{ABC}$.

### 4 Ensemble Modeling

#### 4.1 Deriving model

S100A4 populates both a closed conformation ($M$) and an open conformation ($M^*$), differentiated by exposure of a hydrophobic cleft by rotation of two helices. In the absence of $Ca^{2+}$, $M$ is favored over $M^*$. $Ca^{2+}$ binds cooperatively to four sites in the $M^*$ state [Garrett et al., 2008]. The $M^* \cdot \left(Ca^{2+}\right)_4$ and $M$ species correspond to the "*ca*" and "*apo*" species from the main text. Finally, *peptide* binds preferentially to the $M^*$ state. To model the system, we make the following assumptions:

1. $M$ is strongly favored over $M^*$ in the absence of $Ca^{2+}$.

2. $Ca^{2+}$ binds cooperatively at four equivalent sites on $M^*$.

3. $Ca^{2+}$ binds much more tightly to $M^*$ than $M$, allowing us to neglect the $M \cdot Ca_4^{2+}$ state.

4. *peptide* binds much more tightly to $M^*$ than $M$, allowing us to neglect any $M \cdot peptide$ states.

With these assumptions, we can describe the system with the following scheme and equilibrium constants:

$$M + 4Ca^{2+} \rightleftarrows M^* + 4Ca^{2+} \rightleftarrows M^* \cdot \left(Ca^{2+}\right)_4$$

$$K_* = \frac{[M^*]}{[M]}$$

$$K_C = \frac{\left[M^* \cdot \left(Ca^{2+}\right)_4\right]}{[M^*]\left[Ca^{2+}\right]^4}$$

The stability of $M^* \cdot \left(Ca^{2+}\right)_4$ relative to the other protein conformations is given by:

$$\Delta G = -RTln\left(\frac{\left[M^* \cdot \left(Ca^{2+}\right)_4\right]}{[M] + [M^*]}\right).$$

Substitute the equilibrium constants and simplify:

$$\Delta G = -RTln\left(\frac{[M^*]\, K_C \left[Ca^{2+}\right]^4}{[M] + [M^*]}\right),$$

$$\Delta G = -RTln\left(\frac{K_* [M]\, K_C \left[Ca^{2+}\right]^4}{[M] + K_*[M]}\right),$$

$$\Delta G = -RTln\left(\frac{K_* K_C \left[Ca^{2+}\right]^4}{1+K_*}\right).$$

Assume that $K_* \ll 1$, meaning that $M$ is highly favored over $M^*$ in the absence of $Ca^{2+}$:

$$\Delta G \approx -RTln\left(\frac{K_* K_C \left[Ca^{2+}\right]^4}{1}\right) = -RTln\left(K_* K_C \left[Ca^{2+}\right]^4\right)$$

$$\Delta G = -RTln\left(K_*\right) - RTln\left(K_C\right) - RTln\left(\left[Ca^{2+}\right]^4\right)$$

$$\Delta G = -RTln\left(K_*\right) - RTln\left(K_C\right) - 4RTln\left(\left[Ca^{2+}\right]\right)$$

Setting $\mu_{Ca^{2+}} = RTln([Ca^{2+}])$

$$\Delta G = \Delta G_* + \Delta G_C - 4\mu_{Ca^{2+}}$$

$\Delta G_*$ is the stability of $M^*$ relative to $M$ in the absence of $Ca^{2+}$. $\Delta G_C$ describes the affinity of the $M^*$ state for $Ca^{2+}$. The terms $\Delta G_*$ and $\Delta G_C$, together, describe the intrinsic stability of the active, metal-bound "*ca*" complex at a reference $[Ca^{2+}]$ . We therefore define a new constant:

$$\Delta G^{\circ}_{ca} \equiv \Delta G_* + \Delta G_C$$

The final expression for $\Delta G$ is:

$$\Delta G_{ca}(\mu_{Ca^{2+}}) = \Delta G^{\circ}_{ca} - 4\mu_{Ca^{2+}}$$

The microscopic free energy of the *apo* ($M$) state does not depend on the concentration of $Ca^{2+}$; therefore, $\Delta G_{apo}$ is a constant:

$$\Delta G_{apo}(\mu_{Ca^{2+}}) = \Delta G^{\circ}_{apo}$$

#### 4.2 Setting arbitrary offset

We do not know $\Delta G^{\circ}_{ca}$ or $\Delta G^{\circ}_{apo}$. We do know, however, that at a low calcium concentration $\Delta G_{apo}(\mu_{Ca^{2+}}) \ll \Delta G_{ca}(\mu_{Ca^{2+}})$ (meaning, the $M$ form is favored over $M^*$ at low calcium). We also know that $\Delta G_{ca}(\mu_{Ca^{2+}})$ will increase linearly relative to $\Delta G^{\circ}_{apo}$ as a function of $\mu_{Ca^{2+}}$. If we do not care about the absolute value of $[Ca^{2+}]$ at which the system transitions between favoring *apo* and *pep*, we can choose arbitrary values for $\Delta G^{\circ}_{ca}$ and $\Delta G^{\circ}_{apo}$ and then still calculate how epistasis should change as a function of $\mu_{Ca^{2+}}$ for the protein. For convenience, we set $\Delta G^{\circ}_{apo} = 0$ and $\Delta G^{\circ}_{ca} = 10$ at $\mu_{Ca^{2+}} = 0$.

#### 4.3 Modeling mutant cycles

$ab$ genotype:

$$\Delta G^{ab}_{ca}(\mu_{Ca^{2+}}) = \Delta G^{\circ}_{ca} - 4\mu_{Ca^{2+}}$$

$$\Delta G^{ab}_{apo} = \Delta G^{\circ}_{apo}$$

$$\left\langle \Delta G^{ab}_{apo,ca} \right\rangle (\mu_{Ca^{2+}}) = -RTln\left(e^{-(\Delta G^{\circ}_{ca} - 4\mu_{Ca^{2+}})/RT} + e^{-(\Delta G^{\circ}_{apo})/RT}\right)$$

$Ab$ genotype:

$$\Delta G_{Ab,ca}(\mu_{Ca^{2+}}) = \Delta G^{\circ}_{ca} - 4\mu_{Ca^{2+}} + \Delta\Delta G^{ab}_{A,ca}$$

$$\Delta G_{Ab,apo} = \Delta G^{\circ}_{apo} + \Delta\Delta G^{ab}_{A,apo}$$

$$\left\langle \Delta G^{Ab}_{apo,ca} \right\rangle (\mu_{Ca^{2+}}) = -RTln\left(e^{-(\Delta G^{\circ}_{ca} - 4\mu_{Ca^{2+}} + \Delta\Delta G^{ab}_{A,ca})/RT} + e^{-(\Delta G^{\circ}_{apo} + \Delta\Delta G^{ab}_{A,apo})/RT}\right)$$

$aB$ genotype:

$$\Delta G_{aB,ca}(\mu_{Ca^{2+}}) = \Delta G^{\circ}_{ca} - 4\mu_{Ca^{2+}} + \Delta\Delta G_{B,ca}$$

$$\Delta G_{aB,apo} = \Delta G^{\circ}_{apo} + \Delta\Delta G_{B,apo}$$

$$\left\langle \Delta G^{aB}_{apo,ca} \right\rangle (\mu_{Ca^{2+}}) = -RTln\left(e^{-(\Delta G^{\circ}_{ca} - 4\mu_{Ca^{2+}} + \Delta\Delta G_{B,ca})/RT} + e^{-(\Delta G^{\circ}_{apo} + \Delta\Delta G_{B,apo})/RT}\right)$$

$AB$ genotype:

$$\Delta G_{AB,ca}(\mu_{Ca^{2+}}) = \Delta G^{\circ}_{ca} - 4\mu_{Ca^{2+}} + \Delta\Delta G_{A,ca} + \Delta\Delta G_{B,ca}$$

$$\Delta G_{AB,apo} = \Delta G^{\circ}_{apo} + \Delta\Delta G_{A,apo} + \Delta\Delta G_{B,apo}$$

$$\left\langle \Delta G^{AB}_{apo,ca} \right\rangle (\mu_{Ca^{2+}}) = -RTln\left(e^{-(\Delta G^{\circ}_{ca} - 4\mu_{Ca^{2+}} + \Delta\Delta G_{A,ca} + \Delta\Delta G_{B,ca})/RT} + e^{-(\Delta G^{\circ}_{apo} + \Delta\Delta G_{A,apo} + \Delta\Delta G_{B,apo})/RT}\right)$$

Final expression for $\mu_{Ca^{2+}}$-dependence of $\beta_{AB}$:

$$\beta_{AB}(\mu_{Ca^{2+}}) = \left(\left\langle \Delta G^{Ab}_{ca,apo} \right\rangle - \left\langle \Delta G^{ab}_{ca,apo} \right\rangle\right) + \left(\left\langle \Delta G^{aB}_{ca,apo} \right\rangle - \left\langle \Delta G^{ab}_{ca,apo} \right\rangle\right) - \left(\left\langle \Delta G^{AB}_{ca,apo} \right\rangle - \left\langle \Delta G^{ab}_{ca,apo} \right\rangle\right)$$
